# Malacological survey in a bottle of water: A comparative study between manual sampling and environmental DNA metabarcoding approaches

**DOI:** 10.1101/2020.05.21.108589

**Authors:** Stephen Mulero, Eve Toulza, Anaïs Loisier, Meryl Zimmerman, Jean-François Allienne, Joséphine Foata, Yann Quilichini, Jean-Pierre Pointier, Olivier Rey, Jérôme Boissier

## Abstract

To assess the effect of anthropogenic activities on ecosystems, it is of prime importance to develop new tools enabling a rapid characterisation of ecological communities. Freshwater ecosystems are particularly impacted and threatened by human activities and need thorough attention to preserve their biodiversity and the ecological services they provide. Studying such ecosystems is generally difficult because the associated organisms are hard to sample and to monitor. We present a ready to use environmental metabarcoding diagnostic tool to characterise and monitor the freshwater malacofauna from water samples. The efficiency of this new tool was compared to a classical malacological survey at 19 sampled sites from 10 distinct rivers distributed over Corsica Island (France). Our eDNA monitoring tool demonstrated a remarkable ability to reconstitute the local malacofauna compared to the malacological survey, with 97.1% of species detection confirmed by both methods. The present tool successfully detected the 11 freshwater snail species previously reported in Corsica by malacological survey but was limited at the genus level for some species. Moreover, our malacological survey allowed an update of the local distribution of a wide diversity of freshwater snails including invasive species (i.e. *Potamopyrgus antipodarum* and *Physa acuta*) as well as snail hosts of pathogens of medical and veterinary importance (i.e. *Bulinus truncatus* and *Galba truncatula*).

## Introduction

Anthropogenic activities have adverse effects on ecosystems (Parmesan and Yohe, 2003), including a decrease of biodiversity (Waldron et al., 2017), habitat fragmentation (Strayer and Dudgeon, 2010) and pollution (Blettler et al., 2018). Freshwater ecosystems appear to be particularly threatened by human activities leading to local species extinction (Blettler et al., 2018). According to the IUCN Red list, 46% of these ecosystems worldwide are endangered or vulnerable (Janssen et al., 2016). However, these ecosystems host 10% of all known species despite representing only 0.8% of the Earth’s surface (Strayer and Dudgeon, 2010). Moreover, they also provide important ecological services for the development of human populations (Carpenter et al., 2011).

Over freshwater species, gastropods have received particular attention for several reasons. First, they are suffering massive extinction notably due to habitat loss/degradation, the introduction of alien species and, in some cases, overexploitation (Bouchet et al., 1999, Johnson et al., 2013, Lydeard et al., 2004). Second, they play a key role in the functioning of most freshwater habitats. They are involved in the cycle of several biochemicals especially nitrogen (Hill and Griffiths, 2017) and are at the basis of many food webs especially fishes (Dillon, 2000). Freshwater snails are also reliable bio indicators in particular for detecting the presence of environmental organic or inorganic pollutants such as heavy metals, or microorganisms that are accumulated in their tissues (Mahmoud and Abu Taleb, 2013, Tallarico, 2016). Finally, several freshwater gastropods are of medical or veterinary importance as they constitute intermediate hosts for several pathogens (e.g. *Trematoda*) (Lu et al., 2018). Under the ongoing global changes, these diseases are currently (re-)emerging worldwide at an alarming rate (Kincaid-Smith et al., 2017). Hence, monitoring these snail species is crucial to address risk maps and thus prevent the transmission of these infectious diseases to nearby human and animal populations (Mulero et al., 2020).

Unfortunately, the monitoring of freshwater snails is challenging because: (i) there is no standardised monitoring protocol (Tallarico, 2016); (ii) their populations are spatially and temporally highly dynamic (Lamy et al., 2012); and (iii) classical methods used to collect and to identify snail species are time-consuming, laborious and need malacological expertise (Vinarski and Kramarenko, 2015). In this context, the use of more sensitive molecular detection approaches such as environmental DNA (eDNA) are promising (Bohmann et al., 2014).

Environmental DNA describes the DNA released by living or dead organisms in their environments (Taberlet et al., 2018). This complex and heterogeneous community of DNA fragments originating from all surrounding organisms is either free in the environment or associated with organic or inorganic particles in water, soil or atmosphere (Pedersen et al., 2015, Taberlet et al., 2012). The use of eDNA approaches for monitoring targeted freshwater snail species already exist to detect invasive species such as *Potamopyrgus antipodarum* and *Crepidula fornicata* (Goldberg et al., 2013, Miralles et al., 2019); or gastropod species that transmit parasitic diseases such as *Galba truncatula, Austropeplea tomentosa, Bulinus truncatus* and *Oncomelania hupensis* (Jones et al., 2018, Mulero et al., 2020, Fornillos et al., 2019). In such a biodiversity assessment context, only one eDNA tool have very recently been developed to characterise communities of freshwater bivalves (Prié et al., 2020). However, still is lacking a more general eDNA-based tool to characterise the entire malacofauna.

To fill this gap, we here propose a ready-to-use eDNA-based diagnostic tool for monitoring freshwater snail communities *in natura*. We will first describe our step-by-step procedure used to develop this tool. We will next present a field validation study conducted in the field that was assessed by comparing the snail communities characterised by our eDNA-based diagnostic tool to those obtained by visual malacological survey at 23 sites from 13 rivers widely distributed across Corsica Island (France). Based on our results, we provide an updated vision of the local malacofauna at the Island scale. Finally, we discuss the pros and cons of the emerging eDNA approaches compared to classical prospecting methods.

## Materials and methods

### 1. Malacological survey and water sampling

The field work was conducted during August and September 2018 in the Corsica Island. This summer end period fulfil optimal conditions for the presence of freshwater snails populations that reach their highest density (Kincaid-Smith et al., 2017). Overall a water sampling for eDNA and a malacological survey were performed at 23 sites distributed over 13 different rivers (Fig. 1) and (Table S1). These sites are annually monitored by the French agency “Agence Régionale de Santé” de Corse (ARS, 2019).

**Figure 1.**
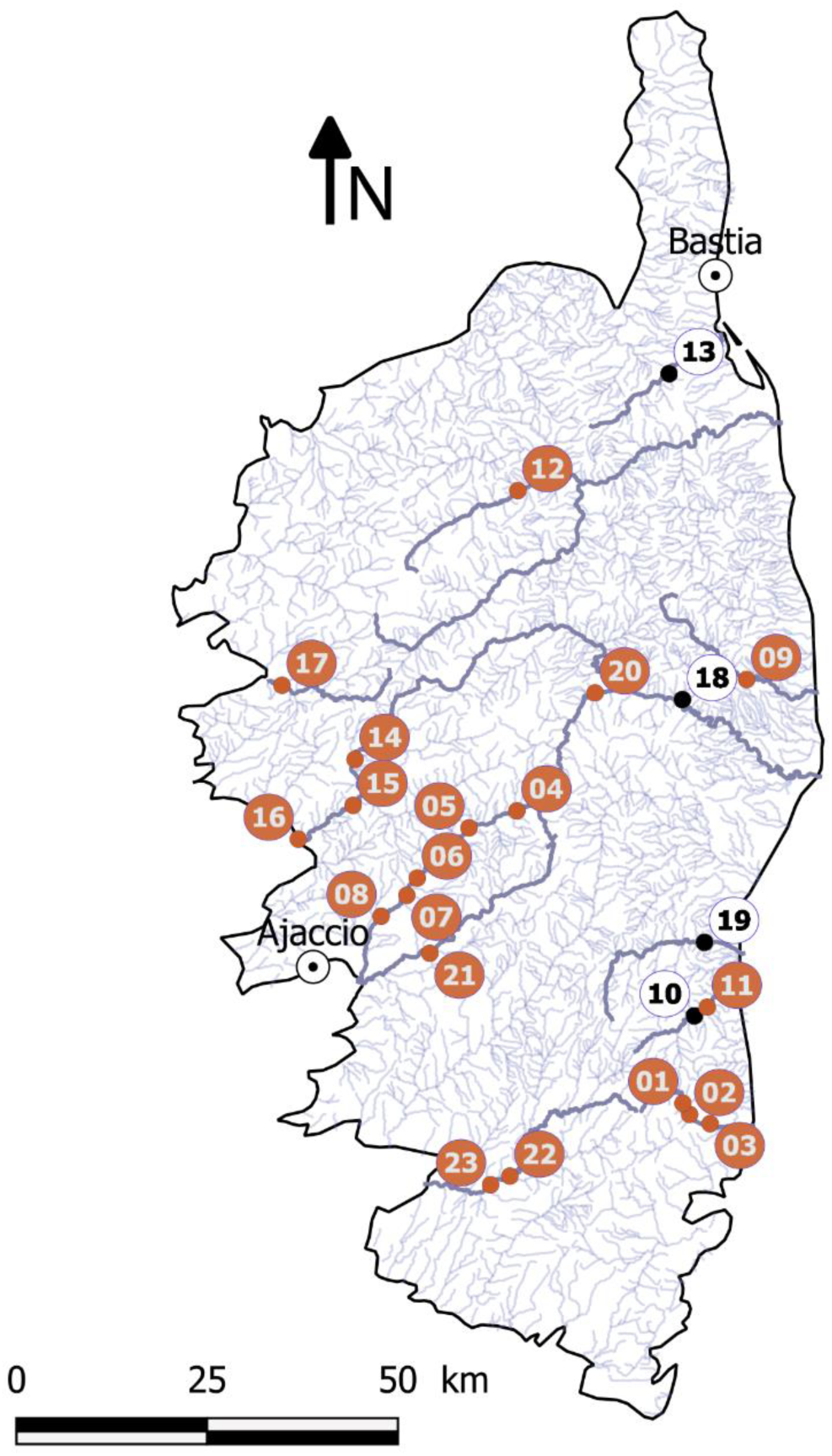
Map referencing the 23 sites prospected across Corsica Island for malacological survey and for water eDNA sampling. All these sites are lotic and distributed among 13 different rivers (darker lines). These sites are annually monitored by the French agency ARS, see (**Table S1**) for coordinates and environmental variables. The sites in black were sampled but removed from the analysis due to potential DNA contaminations (see text for details).

#### 1.1 Water sampling

Water samples for subsequent eDNA analyses were collected before the malacological survey and following a previously optimised protocol (Mulero et al., 2020). At each sampling site, three litres of water were filtered once from the river streambed downstream the prospected site to capture flowing eDNA from upstream transects, covering potentially the overall gastropod communities at a given site, see Supp file 2 for details.

#### 1.2 Malacological survey

At each site, the malacological survey was achieved along a total of 300 meters by the same operators. Three transects of 50 meters (downstream, at the referenced site and upstream) were prospected along the two shores of the river (3 x 2 transects). Within each transect, all freshwater snails were sampled manually or by scooping the grass on the water bench using a colander. Snails were then grouped according to their morphology and taxonomically identified at least at the genus, at best at the species level based on their morphological traits. The number of individuals of each species was semi-quantitatively estimated. If more than 100 snails were collected, we have attributed a “>100” score to the given species. At each site, GPS coordinates and several variables were collected including: the sampling day, the half day of sampling (i.e. A.M. or P.M.), water temperature & pH, water flow and altitude.

### 2. Molecular approaches

All pre-PCR molecular steps were performed under a sterile PCR hood decontaminated before and after each use as follow: the working surface was successively washed using 90% ethanol, 10% bleach and a DNA AWAY^™^ solution (Thermo Scientific^™^) and then exposed to UV light for 30 minutes. The reusable materials were decontaminated following the same protocol as for the field sampling and treated under UV exposure for 30 minutes.

#### 2.1 DNA extractions

Total eDNA from each membrane was extracted using the DNeasy^®^ Blood & Tissue kit (QIAGEN) following an adapted protocol (Mulero et al., 2020). Briefly, membranes were subdivided into four equal parts and extracted following the previously published protocol, see Supp file 2 for details.

In parallel, DNA was extracted from one individual of each of the 11 snail species collected during the malacological survey to produce a synthetic “mock community”. To this aim, we used the E.Z.N.A.^®^ Tissue DNA kit (OMEGA bio-tek, Inc) following the “tissue protocol”. These snail DNA extractions were used as positive PCR and sequencing controls in the following steps.

To refine the molecular species assignations obtained from subsequent NGS analyses, we sequenced a 16S rRNA region for six snail species collected in Corsica. The 16Sbr-H primers were used following the PCR conditions described in (Saito et al., 2018). The resulting amplicons contain the barcode region used for the metabarcoding. The PCR products were then sequenced on an ABI 3730xl sequencer at the GenoScreen platform (Lille, France). Sequences were obtained for *Ancylus fluviatilis* (MT361136), *Gyraulus* sp. (MT361134), *Physa acuta* (MT361133), *Pisidium casertanum* (MT361132), *Theodoxus fluviatilis* (MT361131) and *Galba truncatula* (MT361135).

#### 2.2 PCR primers

The primer pair used in this study (Gast01) was previously developed and tested *in silico* (Taberlet et al., 2018) but never tested and/or applied for empirical purposes. These primers target a 60-70 bp fragment of 16S mitochondrial rRNA. Based on their *in silico* analyses, these primers theoretically amplify at least 1,280 species distributed among 456 genera, mainly related to *Gastropoda*. In our study, we first validated that these primers were able to produce a size-expected amplicon using qPCR for the 11 snail species collected during the malacological survey and eight additional snail species from previous field works (Table 1). These 19 DNA extracts were amplified by qPCR on a LightCycler^®^480 qPCR device following the same protocol as for the first PCR step of the library preparation, see 2.3 below.

**Table 1.** Snail species used for assessing the detection range of the Gast01 PCR primers *in vitro*.

| Snail species | Origin | Date | Provided by | Average headcount per site |
| --- | --- | --- | --- | --- |
| *Bulinus forskalii* | South-Africa | 2017 | Huyse T., KU Leuven, Belgium | – |
| *Bulinus globosus* | Senegal | 2012 | Huyse T., KU Leuven, Belgium | – |
| *Bulinus africanus* | South-Africa | 2017 | Huyse T., KU Leuven, Belgium | – |
| *Radix natalensis*<sup>a,b</sup> | South-Africa | 2017 | Huyse T., KU Leuven, Belgium | – |
| *Galba schirazensis*<sup>a,b</sup> | Cuba | 2013 | Hurtrez-Boussès S., IRD Montpellier, France | – |
| *Galba cubensis*<sup>a,b</sup> | Cuba | 2013 | Hurtrez-Boussès S., IRD Montpellier, France | – |
| *Pseudosuccinea columella*<sup>a,b</sup> | France | 2011 | Hurtrez-Boussès S., IRD Montpellier, France | – |
| *Biomphalaria glabrata*<sup>a,b</sup> | Brazil | 2018 |  | – |
| *Bulinus truncatus*<sup>a,b</sup> | Corsica, France | 2018 | | **2.28** ($\pm 13.95$) from 0 to >100 |
| *Bithynia tentaculata*<sup>b</sup> | Corsica, France | 2018 | | **0.02** ($\pm 0.28$) from 0 to 3 |
| *Radix balthica*<sup>b</sup> | Corsica, France | 2018 | | **0.11** ($\pm 1.21$) from 0 to 13 |
| *Ancylus fluviatilis*<sup>a,b</sup> | Corsica, France | 2018 | | **26.68** ($\pm 40.53$) from 0 to >100 |
| *Galba truncatula*<sup>b</sup> | Corsica, France | 2018 | | **0.06** ($\pm 0.38$) from 0 to 3 |
| *Gyraulus laevis*<sup>a,b</sup> | Corsica, France | 2018 | | **0.06** ($\pm 0.34$) from 0 to 3 |
| *Gyraulus* sp.<sup>a,b</sup> | Corsica, France | 2018 | | **2.99** ($\pm 8.66$) from 0 to 50 |
| *Physa acuta*<sup>a,b</sup> | Corsica, France | 2018 | | **17.38** ($\pm 34.29$) from 0 to >100 |
| *Potamopyrgus antipodarum*<sup>a,b</sup> | Corsica, France | 2018 | | **48.03** ($\pm 46.38$) from 0 to >100 |
| *Pisidium casertanum*<sup>a,b</sup> | Corsica, France | 2018 | | **1.45** ($\pm 7.10$) from 0 to 50 |
| *Theodoxus fluviatilis*<sup>b</sup> | Corsica, France | 2018 | | **0.87** ($\pm 5.26$) from 0 to 50 |

<sup>a</sup>DNA extracts used for the preparation of equimolar pools of DNA (N = 12). <sup>b</sup>DNA extracts used for the preparation of equimolar pools of PCR products (N = 16). The displayed median headcount for each species reported by the malacological survey consider the six measure units conducted on each sites as separate values.

#### 2.3 Library preparation and MiSeq amplicon sequencing

A total of 34 samples including the 23 filtration membranes from the 23 sampling sites, five negative controls and six positive controls were processed. Negative extraction controls consisted in ultrapure water samples processed following the same DNA extraction protocol as for the filtration membranes in each extraction runs (N = 2). PCR negative controls consisted in PCR reactions performed using water as template in each PCR run (N = 3). Positive controls consisted in two categories of mock communities. The first category consisted in an equimolar pool of 12 DNA extracts obtained from individuals of each snail species identified in Corsica and during previous field missions (Table 1). These mock communities are useful to detect PCR competition among different snail species. The second category of mock community consisted in pooling an equimolar quantity of PCR products individually obtained from 16 freshwater snail species (i.e. including the 11 identified snail species collected in Corsica and five additional non-endemic species bred in our laboratory). These controls are useful to detect possible biases during the sequencing process.

Individual libraries were generated for each membrane section of the 23 filtration membranes and for each control, hence, representing a total of 103 libraries (23 x 4 = 92 eDNA samples + 11 controls). NGS libraries were prepared following the Illumina two-step PCR protocol, using mitochondrial 16S primers with Illumina adapters for the first locus-specific PCR. Both PCR were performed using the NEBNext^®^ Ultra^™^ II Q5^®^ Master Mix M0544L (NEW ENGLAND BioLabs, USA). The first PCR reactions were performed in a final volume of 35 µl containing 3.5 µl of extracted DNA (concentrations ranging from 0.05 ng/µl to 8.52 ng/µl), 17.5 µl of 2X Q5 Master Mix and 14 µl of indexed primer mix (final concentration 0.4 µM) and ran on a Techne TC-PLUS thermal cycler PCR device (Techne, UK) using the following program: 98°C for 30 seconds followed by 40 cycles at 98°C for 10 seconds, 55°C for 20 seconds, 65°C for 8 seconds and a final elongation step of 2 minutes at 65°C. At this point, all PCR products were checked on a 2.5% agarose gel stained with ethidium bromide after electrophoresis.

The PCR products were individually indexed in the second PCR step consisting in eight cycles using the Nextera^™^ XT Index (Illumina, San Diego, USA) following the manufacturer’s instructions. Finally, the libraries were normalised using SequalPrep^™^ plates (Thermo Fischer Scientific, USA) before pooling. The pooled libraries were then purified following the JetSeq™ Clean protocol (Bioline, UK), checked on a Bioanalyzer High Sensitivity DNA kit (Agilent, USA) and quantified using a Qubit fluorometric quantification (Thermo Fisher Scientific, USA). Paired-end sequencing (2 x 250 cycles) was performed with a MiSeq Reagent Kit v2 on an Illumina MiSeq^™^ instrument at the Bio-Environnement platform (University of Perpignan Via Domitia, France).

### 3. Data analyses

#### 3.1 Bioinformatic pipeline

The resulting amplicon sequence dataset was processed using the Find Rapidly OTUs with Galaxy Solution (FROGS) pipeline implemented in Galaxy (Escudie et al., 2018) at the Genotoul platform (Toulouse, France). (i) The amplicon dataset was first pre-processed, according to Gast01 primer specificities (i.e. amplicon size of 60 - 70 bp), we filtered out the sequences so as to keep amplicon sizes from 95 to 120 nucleotides (i.e. corresponding to the size of the targeted amplicon with Illumina adapters). (ii) The sequences kept were next clustered in operational taxonomic units (OTUs) using the swarm algorithm and using denoising and an aggregation distance of three nucleotides. (iii) The dataset was filtered out for chimeras using VSEARCH (Rognes et al., 2016). Singletons and underrepresented clusters (i.e. <10 sequences) were removed. (iv) Each OTU was assigned to a species or taxon through a two-step MEGABLAST affiliation procedure. The first MEGABLAST analysis was restricted to *Mollusca* species. The second MEGABLAST analysis was performed without restricting parameters to check the robustness of the previous affiliations. The 10 best hits were kept for subsequent analysis. (v) The resulting OTUs were filtered following two criteria: first, only OTUs presenting a minimal blast coverage of 84% of amplicon length were kept; second, only OTUs presenting a pairwise identity above 89% with the affiliated sequence were kept. The remaining OTUs were considered as “unassigned”.

Lastly, to confirm that a given OTU was detected in a specific sample, we have adapted our validation criteria depending on OTUs abundances. An abundant OTU (i.e. ≥1000 copies considering all the samples) was considered present in a specific membrane when reaching at least 10 copies (overall the four membrane sections). Regarding poorly abundant OTU (i.e. < 1000 copies considering all the samples), we considered a sample positive if four or more copies of such OTU were distributed among two quarters of a same membrane.

## Results

Environmental variables for the 23 sites are presented in (Table S1). All of these sites were lotic ecosystems, pH values ranged from 7.33 to 8.7, temperatures ranged from 16.5°C to 26.7°C and elevation ranged from 4 m to 558 m.

### 1. Malacological survey

Overall, 11 distinct freshwater snail species were identified among the 23 sampled sites (Table 1-S3). These species were already registered in Corsica (INPN, 2020, IUCN, 2020) where 18 species of freshwater mollusc have been described so far, five of which inhabiting lentic ecosystems (Table S3) and (Mouthon, 1982, INPN, 2020). The species richness ranged from one (site 4) to six (site 23). The most common species was *P. antipodarum* (22/23 sites) and the rarest were *T. fluviatilis* (1/23 sites) and *Bithynia tentaculata* (3 individuals at site 13). Among the species identified, two are invasive (i.e. *P. antipodarum* and *P. acuta*) and two are intermediate hosts for Human and livestock parasites (i.e. *B. truncatus* and *G. truncatula*). Our semi-quantitative approach showed that *P. antipodarum* and *A. fluviatilis* were the most abundant species with 48.03±46.38 and 26.68±40.53 individuals, respectively (Table 1).

### 2. Molecular results

Over the 19 snail DNA extracts tested in qPCR with metabarcoding primers (Table 1), all but *P. casertanum* were positive. Regarding the equimolar pools of DNA, whatever pre- (Fig. S4a) or post-PCR (Fig. S4b) the sequencing of equimolar pools gave a number of reads for each species fitting to the expected ratio. This was higher evenness for post-PCR pools.

Despite all our precautions, four among the 23 eDNA water samples (from sites 10, 13, 18 and 19 – see Fig. 1) were possibly contaminated and were thus discarded. In fact, amplicons corresponding of *A. fluviatilis, G. truncatula* and *B. truncatus* were found in the negative PCR controls associated with these samples.

Excluding the four contaminated sites, the whole sequencing (controls + field samples) generated more than 14 M of clusters with an average of 97,000 sequences per library. After analyses, 8.7 M of sequences were affiliated (Fig. 2). These sequences were grouped among 488 OTUs with 43 OTUs corresponding to *Mollusca* species. After removing the sequences from control samples, on the 7,290,067 remaining sequences, 6% (445,638) were affiliated to *Mollusca*; 10% (715,308) were unaffiliated and 84% (6,129,121) were affiliated to non-mollusc species (Fig. 2). Whatever the taxonomic group, the OTU affiliations were generally limited to the genus and rarely reached the species level (e.g. only four out of 11 freshwater snail species found by metabarcoding were affiliated to a single species name). However, the 16S sequences obtained from snail DNA extracts collected in Corsica, allowed to recover the corresponding species name for major *Mollusca* OTUs. All the genus or species detected using eDNA corresponded to species already identified in Corsica and no new genus was detected compared to our malacological survey. Hence, considering all prospected sites, we detected 61.1% (11/18) of historically known freshwater molluscs in Corsica and 84.6% (11/13) of species occurring in lotic systems (Table S3) and (INPN, 2020, Mouthon, 1982). The detected snail species belong to three subclasses of the *Gastropoda* (i.e. *Neritimorpha, Caenogastropoda*, and *Heterobranchia*) (Fig. 3). The non-detected subclasses (e.g. *Vetigastropoda* and *Patellogastropoda*) are related to marine snail species. Beyond gastropods species, the present protocol allowed the detection of the bivalve *P. casertanum*.

**Figure 2.**
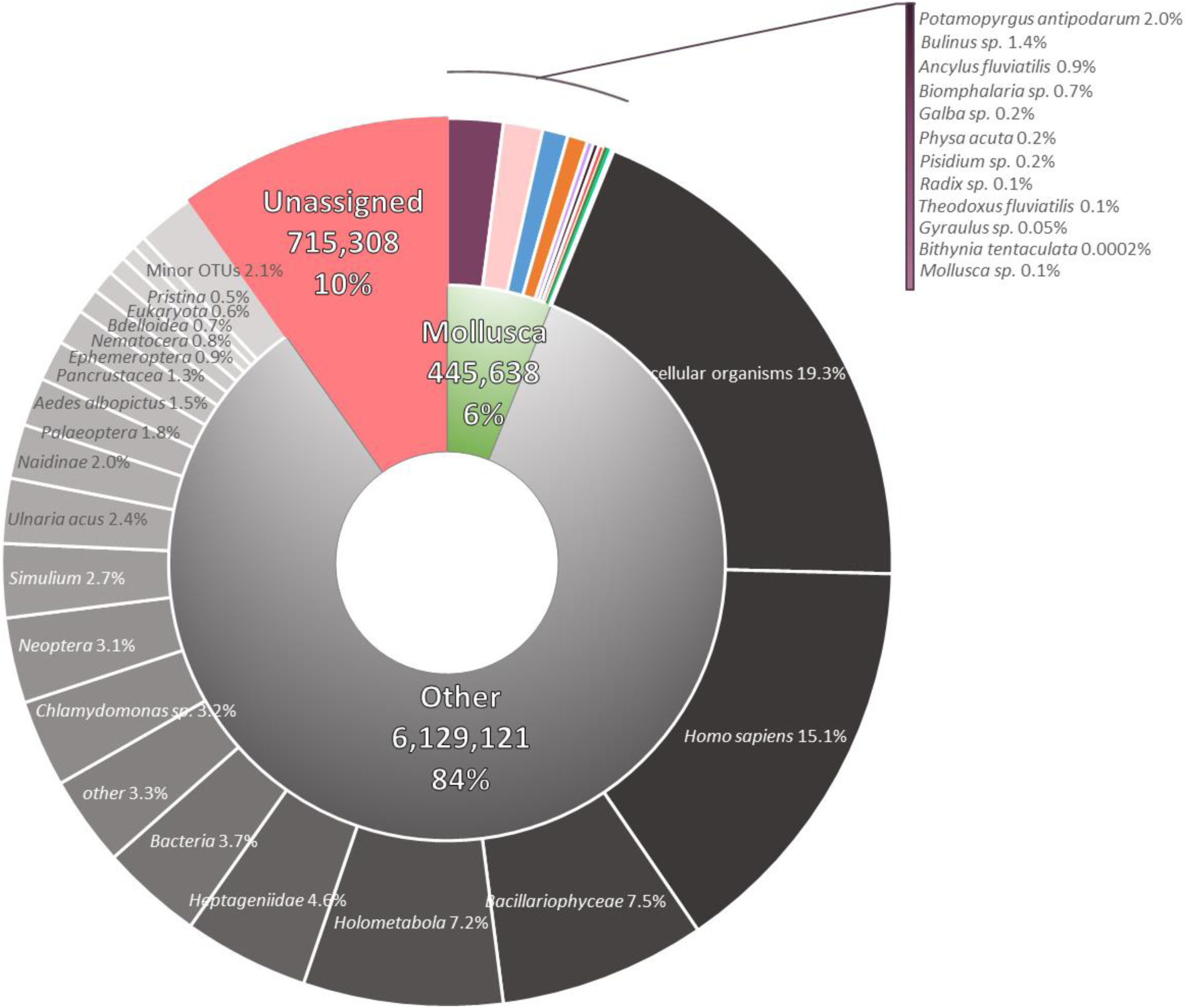
Major OTUs ranking based on sequence abundances. These sequences were obtained from eDNA samples collected in the field and do not contain those from experimental controls. Each taxon is based on merged OTUs (i.e. all OTUs affiliated to a same species are merged in one category).

**Figure 3.**
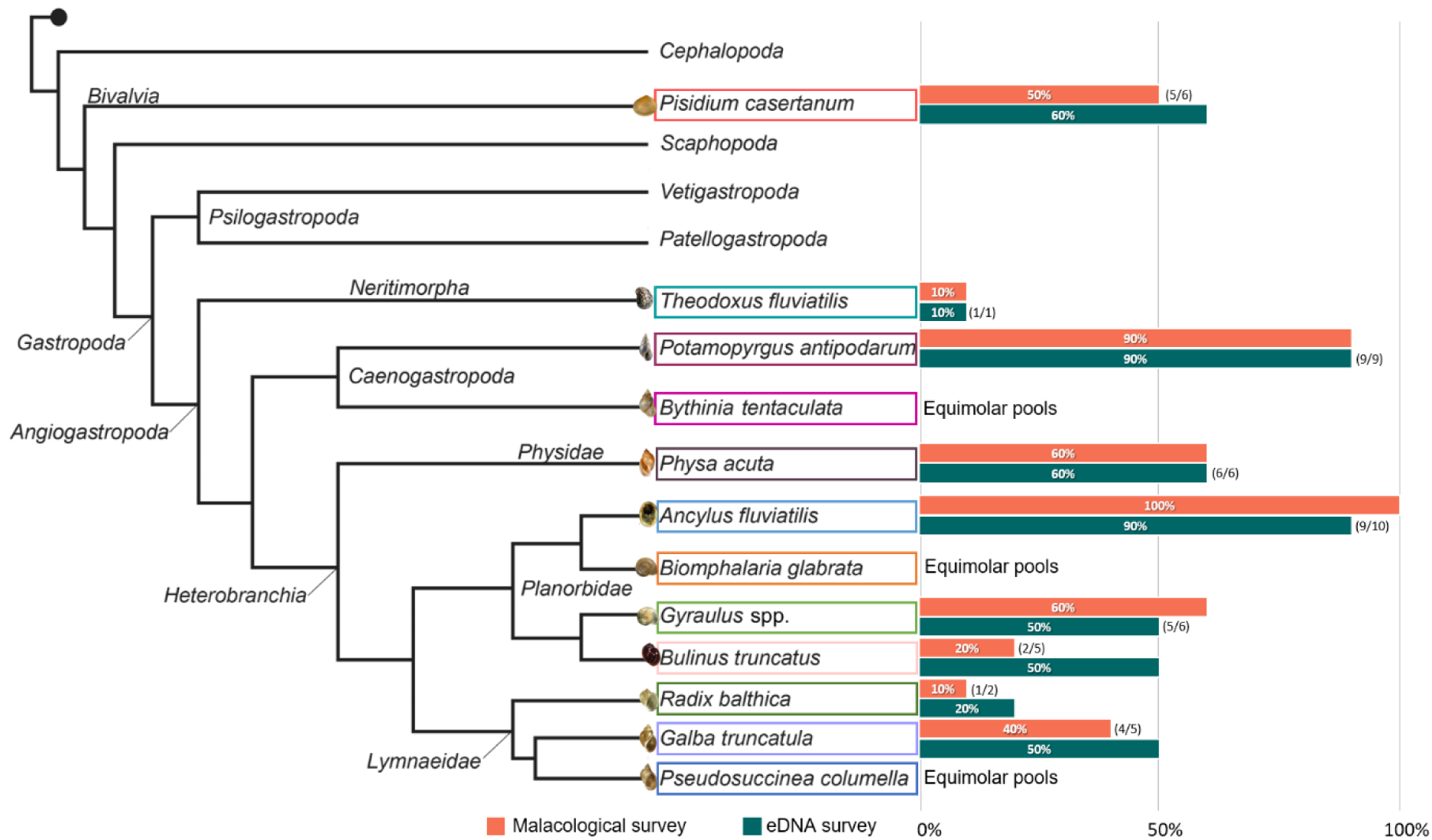
Simplified phylogeny of molluscs from Cunha & Giribet, 2019 highlighting all species reported in Corsica Island during the present study and species used to test primers (mock communities). Horizontal barplots represent species occurrences recorded at the river scale (N = 10) based on the malacological survey (in orange) and on the eDNA metabarcoding results (in blue). Numbers into brackets show the number of rivers positives for both methods on the total of positive rivers for the method showing the highest number of positive for a given species.

### 3. Comparing the malacological survey to the eDNA monitoring

The malacological survey and the eDNA monitoring provided very similar results (Figs 3-4.). At the Island scale, all species identified during the malacological survey were also detected by eDNA monitoring. For some species such as the two *Gyraulus* species, the eDNA monitoring accuracy was limited to the genus level.

At the site level, the eDNA monitoring confirmed the detections obtained by malacological survey for 97.1% (67/69) of the detections. Only two occurrences of *A. fluviatilis* (site 17) and *Gyraulus* sp. (site 20) detected visually during the malacological survey, were not detected by eDNA monitoring (Fig. 4). Conversely, the malacological survey confirmed the detections obtained by eDNA monitoring for 77% (67/87) of the detections (Fig. 4). Moreover, eDNA monitoring detected on average 1.8±2.84 more species per site than the malacological survey although this difference was not significant (Wilcoxon rank test; *W* = 24, *P* = 0.11). Based on the eDNA approach, *B. truncatus* was detected at 11 sites while only visually detected at 4 sites (Figs 4-5a.). At the river scale, the results obtained from direct malacological survey and from the eDNA monitoring approach gave even more congruent results (Fig. 3).

**Figure 4.**
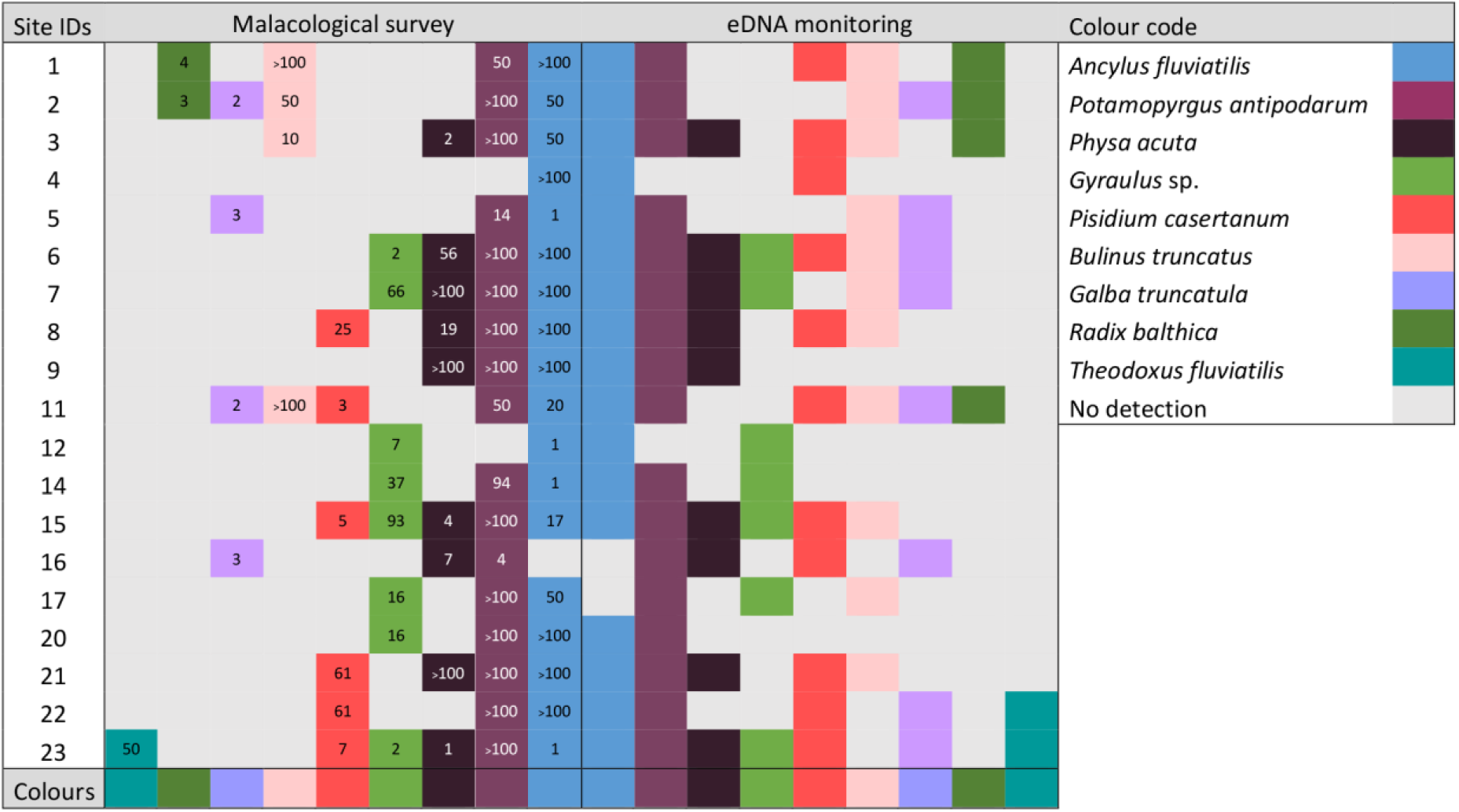
Schematic representation of all detected snail species (or genera) based on the malacological survey (left side) and the eDNA monitoring tool (right side) at the 19 analysed sites (sites 10, 13, 18 and 19 were discarded because of possible DNA contaminations). Columns represent species and are arranged symmetrically between the two detection methods. The number in each cell is the semi-quantitative abundance of each species at each site.

**Figure 5.**
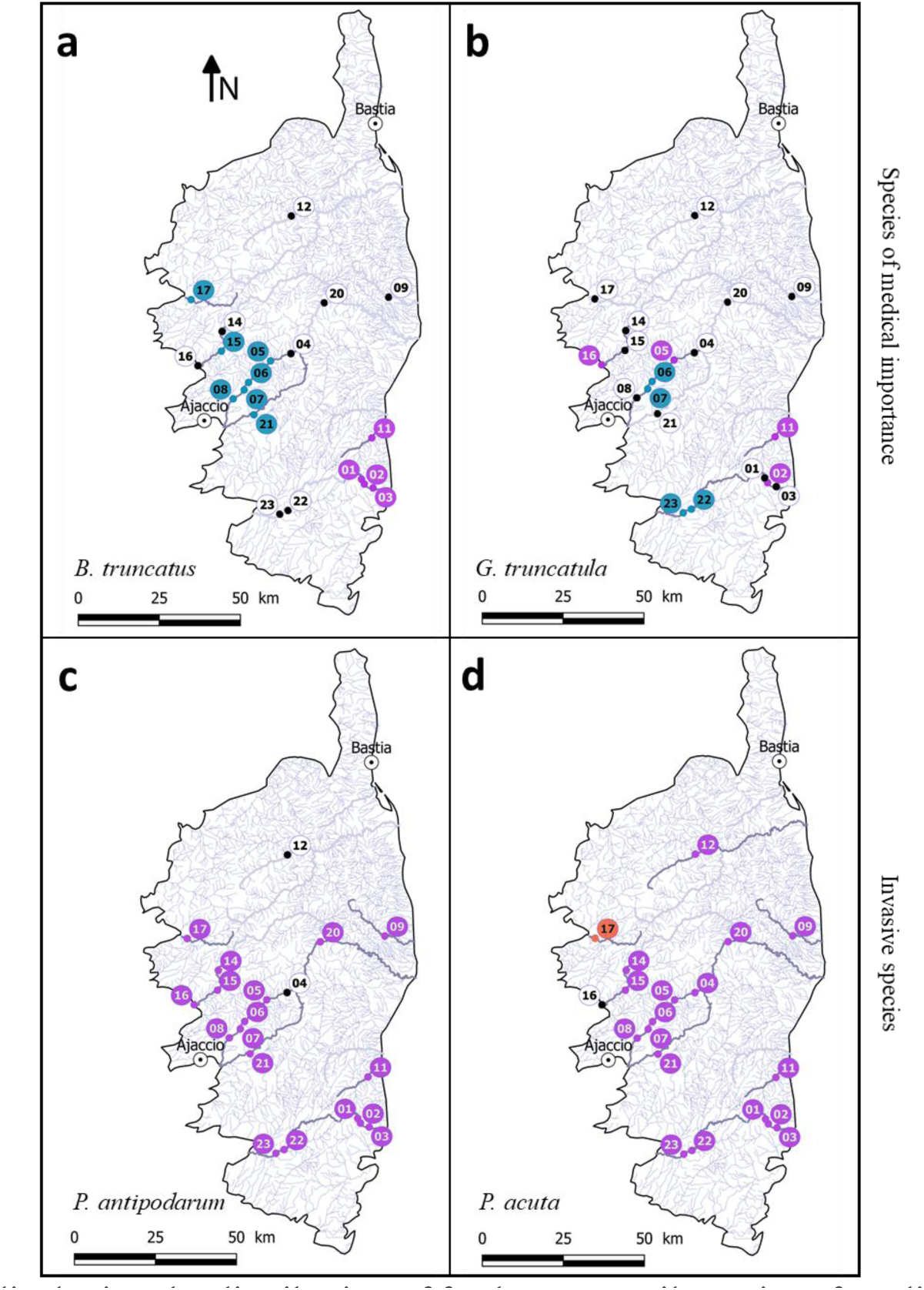
Maps displaying the distribution of freshwater snail species of medical and veterinary (top panels) and ecological interest (down panels), **a.** *Bulinus truncatus*, an intermediate host of the urogenital schistosomiasis infecting Humans; **b.** *Galba truncatula*, an intermediate host of the liver fluke infecting livestock and Humans; **c.** *Potamopyrgus antipodarum* is an invasive snail species from New Zealand and **d.** *Physa acuta* is an invasive snail species from North America. Black dot are sites were the given species is absent based for the two monitoring methods; Purple circles are sites were the given species was detected with the two monitoring methods; Blue circles are sites were the given species was detected using the eDNA monitoring approach only; Orange circles are sites were the given species was detected using the malacological survey only.

Among all the freshwater species detected in Corsica, two species are hosts for the transmission of trematode of medical and veterinary importance (i.e. *B. truncatus*, Fig. 5a and *G. truncatula*, Fig. 5b) and two other are invasive species (i.e. *P. antipodarum*, Fig. 5c and *P. acuta*, Fig. 5d). The two invasive species display a wide distribution range (17 and 9 sites among the 19 monitored sites irrespective of the method) while the two species of medical/veterinary importance were mainly distributed in the Southern part of the Island.

## Discussion

One of the key steps for monitoring ecosystems and to quantify the effect of anthropogenic activities on biodiversity is to develop new tools enabling rapid and exhaustive characterisation of the overall intrinsic species diversity. This is particularly true for freshwater ecosystems that undergo rapid changes and that are particularly threatened nowadays (Carpenter et al., 2011). So far, the use of environmental DNA (eDNA) has been shown to be an efficient tool for monitoring fish (Hanfling et al., 2016), arthropod (Krol et al., 2019) and bivalve (Prié et al., 2020) communities in freshwater environments. Here, we present the first application of an eDNA monitoring method enabling the non-invasive characterisation of entire malacofauna from freshwater samples.

### Benchmarking the eDNA monitoring

Our comparative approach between a classical (i.e. visual) prospection and an eDNA-based malacological survey highlights the remarkable ability of eDNA to characterise malacofauna from water samples. Considering the 19 sites successfully diagnosed, 97.1% (67/69) of total occurrences reported by the malacological survey were confirmed using eDNA monitoring. A noticeable benefit of eDNA approaches compared to malacological survey is the less time-consuming sampling effort needed per site; only one sample of three litres realised by one operator (≈ 30 minutes from sampling to sample preservation) is needed (Mulero et al., 2020). In comparison, the time needed to realise six measure units in classical malacological survey is much longer (>2 hours) depending on field practicability and experimenter skill level. Moreover, provided that the molecular databases are well documented and free of false molecular taxonomic assignations, the eDNA based approach can provide insights on gastropod diversity at a given site without having malacological expertise. In the present study, we did not try to optimise the volume of water sampled. However, in the case of larger sampling campaigns it might be interesting to determine the lowest volume of water necessary to ensure the same diagnosis efficiency. We recently shown, using a targeted approach, that lower water volumes down to one litre allows detecting *B. truncatus* without losses in efficacy (Mulero et al., 2020).

Using the eDNA monitoring approach, we tended to detect more species per site than with the malacological survey. Indeed, over the 87 detection events found using the eDNA monitoring, 77% of them (67) were corroborated by visual observations during the malacological survey. This apparent higher richness found by eDNA metabarcoding approach compared to classical survey was already reported in previous studies (Deiner et al., 2017, Hanfling et al., 2016, Prié et al., 2020). Three non-exclusive hypotheses could explain the higher efficacy of eDNA compare to classical methods. First, DNA transportation through the river flow allows collecting eDNA from upstream the targeted site. In fact, in lotic systems, eDNA is known to disperse in a given way with waterflow (Hanfling et al., 2016). Thus, in lotic systems and depending on the prospected scale, this dispersing property can be favourable for detecting snail communities at a river section scale, but potentially a limiting factor when focusing at the site scale. Second, eDNA-based methods are expected to outcompete visual surveys for detecting cryptic or rare species (e.g. Dejean et al., 2012). Cryptic freshwater mollusc species are particularly difficult to detect and previous studies estimated that about 20% of the species present at a single site are undetected after a single prospecting session during malacological surveys (Dubart et al., 2019). Moreover, detecting biases are also common in malacology because populations of snails are highly dynamic in space and time (Lamy et al., 2012). Interestingly, our eDNA approach has been particularly useful to detect *P. casertanum* compared to classical malacological survey, this species being known to be difficult to detect visually because of his preferred niche (i.e. under boulders) and its small size. Third, false positives constitute one major limitation of eDNA-based protocols (Cristescu and Hebert, 2018). These biases are generally related to either primer amplification biases or inappropriate field and/or laboratory protocols (Cristescu and Hebert, 2018). Regarding primer amplification biases, further applications of the current tool are needed to evaluate these errors considering that it is impossible to test the primers with the DNA of all known organisms *in vitro*. In our study, such biases are likely to be limited because our controlled mock communities were detected with no apparent amplification & sequencing biases between species. Concerning DNA contamination, the four sites that could have been contaminated were discarded, moreover, the detections found using eDNA monitoring yet not confirmed by malacological survey, followed an ecologically realistic distribution. For *G. truncatula*, the two sites found positive only by eDNA monitoring were distributed on the Gravona River, downstream from a site where *G. truncatula* was detected by the two approaches (i.e. site 5). The two other sites were distributed on the Rizzanese river in which *Lymnaeid* snails hosts of *F. hepatica* were previously identified (Gretillat, 1963). Noteworthy, this species prefers temporary water body, but could be found in river shore and associated trickle (Dreyfuss et al., 2009), which might explain the observed relatively low densities.

Regarding *B. truncatus*, the detection differences are the most noticeable between the two approaches, the seven additional sites were mainly distributed in the Gravona River and nearby rivers. Even if these non-randomly distributed detections support rather the non-detection of certain species by malacological survey, than false positive eDNA detection, a thorough field survey of these rivers is needed to validate our results.

### Environmental DNA for characterising communities

The present eDNA metabarcoding protocol allowed to qualitatively characterise the Corsican malacofauna at global scale. However, as shown in previous eDNA-based metabarcoding studies, the accuracy of such characterisation highly depends on the reliability of currently available databases together with sequence size that sometimes results in a limitation for the taxonomic affiliation to the genus taxonomic level especially for closely related species (Krol et al., 2019, Deiner et al., 2017). Here, we couldn’t distinguish the two distinct species belonging to the *Gyraulus* genus identified visually. These species could also be difficult to differentiate morphologically, that could explain confusions with *Planorbis planorbis* on public NCBI databases. At the eDNA era, such limitations clearly call for the improvement and reinforcement of public databases aiming at characterising biodiversity at the taxonomic and molecular scale (Balint et al., 2018). To this end, we submitted 6 new 16S rRNA sequences of freshwater snail species identified in Corsica. However, to overcome these limitations in the immediate future, Krol et al., (2019) recently suggested to use a combination of barcode to optimise taxonomic assignation. Recent studies also called for the need of reinforcing the collection of metadata in eDNA studies that could influence the detection of eDNA (Nicholson et al., 2020). In this way, we collected environmental variables at all sampling sites including some that are generally neglected (e.g. pH, waterflow, water temperature; Nicholson et al., 2020). However, the sampling of a wider diversity of freshwater environment is still needed to accurately assess the effect of these variables on our ability to detect species.

### Environmental DNA for the risk assessment of parasitic diseases

Beyond the technological development, our study provides an updated picture of the distribution of malacological communities in Corsica. Among the 11 species detected, two snail species are of medical/veterinary importance. An updated distribution map for these two species is of prime interest for determining the risk of the associated waterborne diseases. In particular, in 2013 an outbreak of Schistosomiasis emerged in Corsica, this disease being caused by a trematode parasite using *B. truncatus* as host (Boissier et al., 2016, Ramalli et al., 2018). *Bulinus truncatus* was widely distributed in Corsica in the middle of the 20^th^ century and at this time some authors warned authorities about the potential for such a local emergence of Schistosomiasis. Today, this species is mainly established in three rivers in Southern Corsica (Cavu, Solenzara and Osu). Using our eDNA monitoring tool, we detected the presence of *B. truncatus* in the Gravona River, although possibly at low densities and/or upstream the sampled sites. This result call for an urging field validation of the local presence of this species to prevent potential transmission to Humans in this highly frequented river near Ajaccio, the main agglomeration of Corsica.

*Fasciola hepatica* is another trematode species already present in Corsica that uses *G. truncatula* as intermediate host (Oviedo et al., 1996). This parasite is responsible for the zoonotic Fasciolosis disease generally infecting livestock and Humans (Valero et al., 2002). The present study provides an up-to-date geographic distribution of *G. truncatula* in Corsica, hence, delineating pastures presenting a potential risk of transmission to livestock and Humans. Interestingly, the eDNA monitoring tool also detected the presence of *G. truncatula* in the Rizzanese River, a region in which lymnaeids snails hosts of *F. hepatica* were previously identified (Gretillat, 1963) although not locally detected by visual inspection in this study. In a sanitary context, these results also call for thorough monitoring of this species in this particular river.

### Environmental DNA for studying bioinvasions

In a bioinvasion context, we were able to detect two invasive snail species: *P. antipodarum* originating from New Zealand (Goldberg et al., 2013) and introduced in Europe in 1859 probably with human activities such as fish farming and through ballast waters (Alexandre da Silva et al., 2019). This parthenogenetic snail is widely distributed in Corsica and needs attention considering the important density that these populations can reach (>400,000/m^2^). Consequently, this snail species is known to negatively impact nutrient cycling or can competitively exclude native species in invaded ecosystems (Goldberg et al., 2013).

*Physa acuta* is another invasive freshwater snail species originating from North America (Vinarski, 2017) that has invaded Europe likely through the activity of aquarium keepers and maritime trade (Vinarski, 2017). As for *P. antipodarum* this species needs attention as it can drastically disturb the receiving ecosystems and impact the ecology of indigenous species by overconsuming primary producers.

## Authors Contributions

JB., OR. and SM. conceived the study and the sampling design. AL., MZ., JF., YQ and SM. carried out the malacological survey and the field sampling. SM., JFA. and ET. processed the samples and performed molecular experiments and amplicon sequencing. SM., JB., OR. and ET., analysed and interpreted the data. JPP. provided the malacological expertise. SM., JB. and OR. drafted the manuscript. JB., OR., SM., ET., AL., MZ., JFA., JF., YQ. and JPP. revised the manuscript. All authors read and approved the final manuscript.

## Acknowledgements

We thank the French agency “Agence Régionale de Santé de Corse” (ARS) for the data relative to the prospected sites. This study is set within the framework of the “Laboratoire d’Excellence (LABEX)” TULIP (ANR-10-LABX-41).

## Availability of data and materials

The dataset supporting the conclusions of this article is included within the article.

## References

Alexandre Da Silva, M. V., Nunes Souza, J. V., Souza, J. R. B. & Vieira, L. M. 2019. Modelling species distributions to predict areas at risk of invasion by the exotic aquatic New Zealand mudsnail Potamopyrgus antipodarum (Gray 1843). Freshwater Biology, 64, 1504–1518. doi: 10.1111/fwb.13323

ARS. 2019. Qualité des eaux de baignade [Online]. French minister for Solidarity and Health: French minister for Solidarity and Health. Available: http://baignades.sante.gouv.fr/baignades/navigMap.do?idCarte=fra#a [Accessed November 28, 2019 2019].

Balint, M., Nowak, C., Marton, O., Pauls, S. U., Wittwer, C., Aramayo, … Jansen, M. 2018. Accuracy, limitations and cost efficiency of eDNA-based community survey in tropical frogs. Mol Ecol Resour, 18, 1415–1426. doi: 10.1111/1755-0998.12934

Blettler, M. C. M., Abrial, E., Khan, F. R., Sivri, N. & Espinola, L. A. 2018. Freshwater plastic pollution: Recognizing research biases and identifying knowledge gaps. Water Res, 143, 416–424. doi: 10.1016/j.watres.2018.06.015

Bohmann, K., Evans, A., Gilbert, M. T., Carvalho, G. R., Creer, S., Knapp, M., … De Bruyn, M. 2014. Environmental DNA for wildlife biology and biodiversity monitoring. Trends Ecol Evol, 29, 358–67. doi: 10.1016/j.tree.2014.04.003

Boissier, J., Grech-Angelini, S., Webster, B. L., Allienne, J. F., Huyse, T., Mas-Coma, S., … Mitta, G. 2016. Outbreak of urogenital schistosomiasis in Corsica (France): an epidemiological case study. Lancet Infect Dis, 16, 971–9. doi: 10.1016/S1473-3099(16)00175-4

Bouchet, P., Falkner, G. & Seddon, M. B. 1999. Lists of protected land and freshwater molluscs in the Bern Convention and European Habitats Directive: are they relevant to conservation? Biological Conservation, 90, 21–31.

Carpenter, S. R., Stanley, E. H. & Vander Zanden, M. J. 2011. State of the World’s Freshwater Ecosystems: Physical, Chemical, and Biological Changes. Annual Review of Environment and Resources, 36, 75–99. doi: 10.1146/annurev-environ-021810-094524

Cristescu, M. E. & Hebert, P. D. N. 2018. Uses and Misuses of Environmental DNA in Biodiversity Science and Conservation. Annual Review of Ecology, Evolution, and Systematics, 49, 209–230. doi: 10.1146/annurev-ecolsys-110617-062306

Cunha, T. J. & Giribet, G. 2019. A congruent topology for deep gastropod relationships. Proc Biol Sci, 286, 20182776. doi: 10.1098/rspb.2018.2776

Deiner, K., Bik, H. M., Machler, E., Seymour, M., Lacoursiere-Roussel, A., Altermatt, F., Creer, S., … & Bernatchez, L. 2017. Environmental DNA metabarcoding: Transforming how we survey animal and plant communities. Mol Ecol, 26, 5872–5895. doi: 10.1111/mec.14350

Dejean, T., Valentini, A., Miquel, C., Taberlet, P., Bellemain, E. & Miaud, C. 2012. Improved detection of an alien invasive species through environmental DNA barcoding: the example of the American bullfrog *Lithobates catesbeianus*. Journal of Applied Ecology, 49, 953–959. doi: 10.1111/j.1365-2664.2012.02171.x

Dillon, R. T. 2000. Predation. The Ecology of Freshwater Molluscs. Cambridge: Cambridge University Press. doi: 10.1017/cbo9780511542008.008

Dreyfuss, G., Vareille-Morel, V. & Rondelaud, D. 2009. Les habitats de *Lymnaea truncatula* Müller (Mollusque) le long de deux rivières. Annales de Limnologie - International Journal of Limnology, 33, 67–72. doi: 10.1051/limn/1997009

Dubart, M., Pantel, J. H., Pointier, J. P., Jarne, P. & David, P. 2019. Modeling competition, niche, and coexistence between an invasive and a native species in a two-species metapopulation. Ecology, 100, e02700. doi: 10.1002/ecy.2700

Escudie, F., Auer, L., Bernard, M., Mariadassou, M., Cauquil, L., Vidal, K., … Pascal, G. 2018. FROGS: Find, Rapidly, OTUs with Galaxy Solution. Bioinformatics, 34, 1287–1294. doi: 10.1093/bioinformatics/btx791

Fornillos, R. J. C., Sato, M. O., Tabios, I. K. B., Sato, M., Leonardo, L. R., Chigusa, Y., … Fontanilla, I. K. C. 2019. Detection of *Schistosoma japonicum* and *Oncomelania hupensis quadrasi* environmental DNA and its potential utility to schistosomiasis japonica surveillance in the Philippines. PLoS One, 14, e0224617. doi: 10.1371/journal.pone.0224617

Goldberg, C. S., Sepulveda, A., Ray, A., Baumgardt, J. & Waits, L. P. 2013. Environmental DNA as a new method for early detection of New Zealand mudsnails (*Potamopyrgus antipodarum*). Freshwater Science, 32, 792–800. doi: 10.1899/13-046.1

Gretillat, S. 1963. [Epidemiology of Certain Trematode Diseases of Domestic Animals in Corsica (Bovine Bilharziosis and Bovine and Ovine Distomiasis). Observations Conducted during a Mission Accomplished during the Autumn of 1962]. Ann Parasitol Hum Comp, 38, 471–81.

Hanfling, B., Lawson Handley, L., Read, D. S., Hahn, C., Li, J., Nichols, P., Blackman, R. C., … Winfield, I. J. 2016. Environmental DNA metabarcoding of lake fish communities reflects long-term data from established survey methods. Mol Ecol, 25, 3101–19. doi: 10.1111/mec.13660

Hill, W. R. & Griffiths, N. A. 2017. Nitrogen processing by grazers in a headwater stream: riparian connections. Freshwater Biology, 62, 17–29. doi: 10.1111/fwb.12833

INPN. 2020. National Inventory of Natural Heritage [Online]. Muséum national d’Histoire naturelle. Available: https://inpn.mnhn.fr [Accessed February 11 2020].

IUCN. 2020. International Union for Conservation of Nature [Online]. International Union for Conservation of Nature. Available: www.iucn.org [Accessed February 10 2020].

Janssen, J. A. M., Rodwell, J. S., Garcia-Criado, S., Gubbay, S., Haynes, T., Nieto, A., … Valachovic, M. 2016. European Red List of Habitats Part 2. Terrestrial and freshwater habitats. European Publications Office, 2. doi: 10.2779/091372

Johnson, P. D., Bogan, A. E., Brown, K. M., Burkhead, N. M., Cordeiro, J. R., Garner, J. T., … Strong, E. E. 2013. Conservation status of freshwater gastropods of Canada and the United States. Fisheries, 38, 247–282.

Jones, R. A., Brophy, P. M., Davis, C. N., Davies, T. E., Emberson, H., Rees Stevens, P. & Williams, H. W. 2018. Detection of *Galba truncatula, Fasciola hepatica* and *Calicophoron daubneyi* environmental DNA within water sources on pasture land, a future tool for fluke control? Parasit Vectors, 11, 342. doi: 10.1186/s13071-018-2928-z

Kier, G., Kreft, H., Lee, T. M., Jetz, W., Ibisch, P. L., Nowicki, C., … Barthlott, W. 2009. A global assessment of endemism and species richness across island and mainland regions. Proc Natl Acad Sci U S A, 106, 9322–7. doi: 10.1073/pnas.0810306106

Kincaid-Smith, J., Rey, O., Toulza, E., Berry, A. & Boissier, J. 2017. Emerging Schistosomiasis in Europe: A Need to Quantify the Risks. Trends Parasitol, 33, 600–609. doi: 10.1016/j.pt.2017.04.009

Krol, L., Van Der Hoorn, B., Gorsich, E. E., Trimbos, K., Bodegom, P. M. V. & Schrama, M. 2019. How Does eDNA Compare to Traditional Trapping? Detecting Mosquito Communities in South-African Freshwater Ponds. Frontiers in Ecology and Evolution, 7. doi: 10.3389/fevo.2019.00260

Lamy, T., Pointier, J. P., Jarne, P. & David, P. 2012. Testing metapopulation dynamics using genetic, demographic and ecological data. Mol Ecol, 21, 1394–410. doi: 10.1111/j.1365-294X.2012.05478.x

Longmire, J. L., Maltbie, M. & Baker, R. J. 1997. Use of “lysis buffer” in DNA isolation and its implication for museum collections. Museum of Texas Tech University.

Lu, X. T., Gu, Q. Y., Limpanont, Y., Song, L. G., Wu, Z. D., Okanurak, K. & Lv, Z. Y. 2018. Snail-borne parasitic diseases: an update on global epidemiological distribution, transmission interruption and control methods. Infect Dis Poverty, 7, 28. doi: 10.1186/s40249-018-0414-7

Lydeard, C., Cowie, R. H., Ponder, W. F., Bogan, A. E., Bouchet, P., Clark, S. A., … Thompson, F. G. 2004. The global decline of nonmarine mollusks. BioScience, 54, 321–330.

Mahmoud, K. M. A. & Abu Taleb, H. M. A. 2013. Fresh water snails as bioindicator for some heavy metals in the aquatic environment. African Journal of Ecology, 51, 193–198.

Miralles, L., Parrondo, M., Hernandez De Rojas, A., Garcia-Vazquez, E. & Borrell, Y. J. 2019. Development and validation of eDNA markers for the detection of *Crepidula fornicata* in environmental samples. Mar Pollut Bull, 146, 827–830. doi: 10.1016/j.marpolbul.2019.07.050

Mouthon, J. 1982. Les mollusques dulcicoles - Données biologiques et écologiques - Clés de détermination des principaux genres de bivalves et de gastéropodes de France. Bulletin Français de Pisciculture, 1–27. doi: 10.1051/kmae:1982001

Mulero, S., Boissier, J., Allienne, J., Quilichini, Y., Foata, J., Pointier, J. & Rey, O. 2020. Environmental DNA for detecting *Bulinus truncatus*: A new environmental surveillance tool for schistosomiasis emergence risk assessment. Environmental DNA, 2, 161–174. doi: 10.1002/edn3.53

Nicholson, A., Mcisaac, D., Macdonald, C., Gec, P., Mason, B. E., Rein, W., … Hanner, R. H. 2020. An analysis of metadata reporting in freshwater environmental DNA research calls for the development of best practice guidelines. Environmental DNA. doi: 10.1002/edn3.81.

Oviedo, J. A., Bargues, M. D. & Mas-Coma, S. 1996. The intermediate snail host of *Fasciola hepatica* on the mediterranean island of Corsica. A.P.E., 56, 217–220.

Parmesan, C. & Yohe, G. 2003. A globally coherent fingerprint of climate change impacts across natural systems. Nature, 421, 37–42. doi: 10.1038/nature01286

Pedersen, M. W., Overballe-Petersen, S., Ermini, L., Sarkissian, C. D., Haile, J., Hellstrom, M., … Willerslev, E. 2015. Ancient and modern environmental DNA. Philos Trans R Soc Lond B Biol Sci, 370, 20130383. doi: 10.1098/rstb.2013.0383

Prié, V., Valentini, A., Lopes-Lima, M., Froufe, E., Rocle, M., Poulet, N., Taberlet, P. & Dejean, T. 2020. Environmental DNA metabarcoding for freshwater bivalves biodiversity assessment: methods and results for the Western Palearctic (European sub-region). Hydrobiologia. doi: 10.1007/s10750-020-04260-8.

Ramalli, L., Mulero, S., Noel, H., Chiappini, J. D., Vincent, J., Barre-Cardi, H., … Boissier, J. & Berry, A. 2018. Persistence of schistosomal transmission linked to the Cavu river in southern Corsica since 2013. Euro Surveill, 23, 2–5. doi: 10.2807/1560-7917.ES.2018.23.4.18-00017

Rognes, T., Flouri, T., Nichols, B., Quince, C. & Mahe, F. 2016. VSEARCH: a versatile open source tool for metagenomics. PeerJ, 4, e2584. doi: 10.7717/peerj.2584

Saito, T., Hirano, T., Prozorova, L., Tu Do, V., Sulikowska-Drozd, A., Sitnikova, T., … Fukuda, H. & Chiba, S. 2018. Phylogeography of freshwater planorbid snails reveals diversification patterns in Eurasian continental islands. BMC Evol Biol, 18, 164. doi: 10.1186/s12862-018-1273-3

Strayer, D. L. & Dudgeon, D. 2010. Freshwater biodiversity conservation: recent progress and future challenges. Journal of the North American Benthological Society, 29, 344–358. doi: 10.1899/08-171.1

Taberlet, P., Bonin, A., Zinger, L. & Coissac, E. 2018. Environmental DNA: For biodiversity research and monitoring, Oxford, UK, Oxford University Press.

Taberlet, P., Coissac, E., Hajibabaei, M. & Rieseberg, L. H. 2012. Environmental DNA. Mol Ecol, 21, 1789–93. doi: 10.1111/j.1365-294X.2012.05542.x

Tallarico, L. D. F. 2016. Freshwater Gastropods as a Tool for Ecotoxicology Assessments in Latin America*. American Malacological Bulletin, 33, 330–336. doi: 10.4003/006.033.0220

Valero, M. A., Panova, M., Comes, A. M., Fons, R. & Mas-Coma, S. 2002. Patterns in size and shedding of *Fasciola hepatica* eggs by naturally and experimentally infected murid rodents. J Parasitol, 88, 308–13. doi: 10.1645/0022-3395(2002)088[0308:PISASO]2.0.CO;2

Vinarski, M. V. 2017. The history of an invasion: phases of the explosive spread of the physid snail *Physella acuta* through Europe, Transcaucasia and Central Asia. Biological Invasions, 19, 1299–1314. doi: 10.1007/s10530-016-1339-3

Vinarski, M. V. & Kramarenko, S. S. 2015. How does the discrepancies among taxonomists affect macroecological patterns? A case study of freshwater snails of Western Siberia. Biodiversity and Conservation, 24, 2079–2091. doi: 10.1007/s10531-015-0934-4

Waldron, A., Miller, D. C., Redding, D., Mooers, A., Kuhn, T. S., Nibbelink, N., Roberts, J. T., Tobias, J. A. & Gittleman, J. L. 2017. Reductions in global biodiversity loss predicted from conservation spending. Nature, 551, 364–367. doi: 10.1038/nature24295

