## Supplementary material for "Malacological survey in a bottle of water: A comparative study between manual sampling and environmental DNA metabarcoding approaches": S1

**Table S1.** Environmental variables and geographical coordinates (using WGS 84 projection) for each of the prospected sites.

| **Site IDs** | **River** | **Coordinates (DD)** | | **Sampling Date** | **Water flow** | **T (°C)** | **pH** | **Altitude (m)** |
| --- | --- | --- | --- | --- | --- | --- | --- | --- |
| 1 | Cavu | 41.731388 | 9.293333 | 26/09/18 - am | low to strong | 18.8 | 8.7 | 157 |
| 2 | Cavu | 41.72 | 9.301666 | 26/09/18 - am | medium | 20 | 8.5 | 124 |
| 3 | Cavu | 41.704444 | 9.334722 | 26/09/18 - pm | strong | 21.5 | 8.3 | 61 |
| 4 | Gravona | 42.093265 | 9.067898 | 20/08/18 - am | low | 16.5 | 7.41 | 558 |
| 5 | Gravona | 42.077132 | 8.99049 | 20/08/18 - am | low to strong | 17.3 | 7.6 | 341 |
| 6 | Gravona | 42.003029 | 8.884666 | 17/08/18 - am | low to medium | 20.8 | 7.95 | 113 |
| 7 | Gravona | 42.023167 | 8.903336 | 17/08/18 - pm | none to low | 21.6 | 7.96 | 172 |
| 8 | Gravona | 41.981789 | 8.841893 | 23/08/18 - am | low to strong | 20.9 | 7.87 | 65 |
| 9 | La Bravone | 42.225705 | 9.446491 | 04/08/18 - am | low | 22.5 | 8.31 | 84 |
| 10 | La Solenzara | 41.8348 | 9.319537 | 31/07/18 - pm | low | 26.7 | 7.68 | 114 |
| 11 | La Solenzara | 41.843611 | 9.343333 | 27/09/18 - am | low to strong | 19.5 | 8.3 | 62 |
| 12 | L'Asco | 42.467624 | 9.10532 | 07/08/18 - am | none to low | 22.8 | 7.56 | 351 |
| 13 | Le Bevinco | 42.591671 | 9.36287 | 03/08/18 - am | none to low | 20.7 | 8.21 | 318 |
| 14 | Le Liamone | 42.168943 | 8.818982 | 13/08/18 - pm | none to medium | 23.6 | 7.89 | 184 |
| 15 | Le Liamone | 42.078413 | 8.720191 | 15/08/18 - am | none to low | 23.3 | 7.33 | 4 |
| 16 | Le Liamone | 42.116121 | 8.817636 | 15/08/18 - am | low to strong | 21.2 | 7.89 | 50 |
| 17 | Le Porto | 42.260545 | 8.712663 | 13/08/18 - am | none to medium | 21.9 | 7.92 | 32 |
| 18 | Le Tavignano | 42.209283 | 9.343659 | 01/08/18 - am | low to medium | 24.7 | 8.14 | 129 |
| 19 | Le Travo | 41.920442 | 9.346367 | 31/07/18 - am | low to medium | 23.7 | 7.89 | 30 |
| 20 | Le Vecchio | 42.224401 | 9.203844 | 06/08/18 - am | low | 22.1 | 7.74 | 223 |
| 21 | Prunelli | 41.932864 | 8.914213 | 23/08/18 - am | low to strong | 17.9 | 7.62 | 109 |
| 22 | Rizzanese | 41.664014 | 9.013416 | 21/08/18 - am | none to strong | 20 | 7.9 | 39 |
| 23 | Rizzanese | 41.655436 | 8.982305 | 21/08/18 - am | none to medium | 21.4 | 8.22 | 48 |

The water flow index (m/s) was measured along the prospected transect on the water surface with a floating object, strong (≥1 m/s), medium (0.5 m/s), low (0.1 m/s) and none (0 m/s).
