## Supplementary material for "Malacological survey in a bottle of water: A comparative study between manual sampling and environmental DNA metabarcoding approaches": Supp file 2

**Supplementary file 2**: Details on water sampling protocol and DNA extraction protocol.

**Water Sampling**

Water filtrations were achieved using a sterile filtration unit with a 0.45 µm PES membrane of 90 mm diameter (VWR PES filter unit 1 L, model 514-0301) connected to a manual vacuum pump (Mityvac model MV8500). Afterwards, each filtration membrane was immediately stored in a 50 mL tube with a sterile Longmire buffer (Longmire et al., 1997). Membranes were then stored in a dark cold room at 4°C until subsequent molecular analyses. During all the sampling process, precautions were taken to avoid any contamination (Taberlet et al., 2012): operators used disposable sterile gloves and all reusable materials (i.e. scissors and forceps) were decontaminated before and after each site by successively placing instruments in a 10% bleach bath, in a 90% ethanol bath and then in a DNA AWAY^™^ bath (Thermo Scientific) finalised by a flame sterilisation (Taberlet et al., 2012).

**Environmental DNA extraction**

Membranes were subdivided into four equal parts to be lysed in 2 ml microcentrifuge tubes at 65°C for one hour in a solution containing 567 µl of ATL buffers and 63 µl of proteinase K solution. Next, 630 µl of AL buffers and 630 µl of 100% ethanol were added. The remaining steps were performed following the manufacturer instructions, and DNA was ultimately eluted in 100 µl (2 * 50 µl) of AE buffer pre-heated to 65°C. The resulting DNA extracts were then preserved at -20°C until subsequent steps.
