## Supplementary material for "Malacological survey in a bottle of water: A comparative study between manual sampling and environmental DNA metabarcoding approaches": S3

**Table S3.** Freshwater molluscs that were historically and presently detected in Corsica.

| **Species name** | **Habitat** |  | **Identified in sites** | **Average headcount per site** |
| --- | --- | --- | --- | --- |
| *Galba truncatula* | Lotic, Lentic |  | 2, 5, 11, 16 | **0.06** (± 0.38) from 0 to 3 |
| *Bulinus truncatus* | Lotic, Lentic |  | 1, 2, 3, 11 | **2.28** (± 13.95) from 0 to >100 |
| *Bithynia tentaculata* | Lotic, Lentic |  | 13 | **0.02** (± 0.28) from 0 to 3 |
| *Ancylus fluviatilis* | Lotic |  | All except 16 | **26.68** (± 40.53) from 0 to >100 |
| *Potamopyrgus antipodarum* | Lotic, Lentic |  | All except 4 and 12 | **48.03** (± 46.38) from 0 to >100 |
| *Physa acuta* | Lentic, Lotic |  | 3, 6, 7, 8, 9, 15, 16, 21, 23 | **17.38** (± 34.29) from 0 to >100 |
| *Pisidium casertanum* | Lotic, ditches, ponds |  | 8, 11, 15, 21, 22, 23 | **1.45** (± 7.10) from 0 to 50 |
| *Gyraulus laevis* | Lentic, lotic |  | 20, 23 | **0.06** (± 0.34) from 0 to 3 |
| *Gyraulus* sp. | Lentic, lotic |  | 6, 7, 12, 14, 15, 17, 20, 23 | **2.99** (± 8.66) from 0 to 50 |
| *Radix balthica* | Lentic, Lotic |  | 1, 2 | **0.11** (± 1.21) from 0 to 13 |
| *Theodoxus fluviatilis* | Lotic |  | 23 | **0.87** (± 5.26) from 0 to 50 |
| *Stagnicola palustris* | Ponds |  | _ |  |
| *Planorbis planorbis* | Ponds, lentic |  | _ |  |
| *Moitessieria corsica* | Groundwater |  | _ |  |
| *Sphaerium lacustre* | Lentic |  | _ |  |
| *Unio mancus* | Lotic |  | _ |  |
| *Acroloxus lacustris* | Lentic |  | _ |  |
| *Hippeutis complanatus* | Ponds, ditches |  | _ |  |

These data are recovered from the work of Mouthon, 1982 and confirmed with the INPN 2020 database. Lentic = still water, Lotic = flowing water. The displayed median headcount for each species reported by the malacological survey consider the six measure units conducted on each sites as separate values.
