## Supplementary material for "Malacological survey in a bottle of water: A comparative study between manual sampling and environmental DNA metabarcoding approaches": Fig. S4

**b**

**a**

|  | Ancylus sp. fluviatilis | Biomphalaria sp. | Bithynia tentaculata | Bulinus sp. | Galba sp. | Gyraulus sp. | Physa acuta | Pisidium sp. | Potamopyrgus antipodarum | Pseudosuccinea columella | Theodoxus fluviatilis |
| --- | --- | --- | --- | --- | --- | --- | --- | --- | --- | --- | --- |
| Rep 1 | 20749 | 27390 | 8 | 5854 | 41045 | 34137 | 10481 | 25502 | 10752 | 4018 | 9 |
| Rep 2 | 19773 | 26584 | 9 | 5830 | 36241 | 32774 | 8765 | 22568 | 9680 | 3409 | 11 |
| Rep 3 | 17611 | 23368 | 13 | 4834 | 31574 | 28749 | 7038 | 21032 | 8385 | 3000 | 12 |
| Expected | 1 | 1 | 0 | 1 | 2 | 2 | 1 | 1 | 1 | 1 | 1 |

|  | Ancylus sp. fluviatilis | Biomphalaria sp. | Bithynia tentaculata | Bulinus sp. | Galba sp. | Gyraulus sp. | Physa acuta | Pisidium sp. | Potamopyrgus antipodarum | Pseudosuccinea columella | Radix sp. | Theodoxus fluviatilis |
| --- | --- | --- | --- | --- | --- | --- | --- | --- | --- | --- | --- | --- |
| Rep 1 | 12609 | 22685 | 7495 | 10591 | 26244 | 12208 | 15499 | 15850 | 12393 | 11515 | 18335 | 7395 |
| Rep 2 | 9476 | 17109 | 5551 | 7816 | 19263 | 9086 | 11901 | 13083 | 9553 | 9043 | 14222 | 6294 |
| Rep 3 | 10078 | 17895 | 6123 | 8254 | 20256 | 9591 | 12233 | 13944 | 10101 | 9523 | 15033 | 6530 |
| Expected | 1 | 1 | 1 | 1 | 3 | 2 | 1 | 1 | 1 | 1 | 2 | 1 |

**Figure S4.** Number of sequences generated per species in equimolar controls, **a**. results for pooled DNA (N = 12) and **b.** results for pooled PCR products (N = 16). The first three columns are associated to the three technical replicate realised for each pool category, the fourth column give the expected composition of the sample in number of distinct DNA ordered per genus.
